# Space-by-time modular decomposition effectively describes whole-body muscle activity during upright reaching in various directions

**DOI:** 10.1101/155085

**Authors:** Pauline M. Hilt, Ioannis Delis, Thierry Pozzo, Bastien Berret

## Abstract

The modular control hypothesis suggests that motor commands are built from precoded modules whose specific combined recruitment can allow the performance of virtually any motor task. Despite considerable experimental support, this hypothesis remains tentative as classical findings of reduced dimensionality in muscle activity may also result from other constraints (biomechanical couplings, data averaging or low dimensionality of motor tasks). Here we assessed the effectiveness of modularity in describing muscle activity in a comprehensive experiment comprising 72 distinct point-to-point whole-body movements during which the activity of 30 muscles was recorded. To identify invariant modules of a temporal and spatial nature, we used a space-by-time decomposition of muscle activity that has been shown to encompass classical modularity models. To examine the decompositions, we focused not only on the amount of variance they explained but also on whether the task performed on each trial could be decoded from the single-trial activations of modules. For the sake of comparison, we confronted these scores to the scores obtained from alternative non-modular descriptions of the muscle data. We found that the space-by-time decomposition was effective in terms of data approximation and task discrimination at comparable reduction of dimensionality. These findings show that few spatial and temporal modules give a compact yet approximate representation of muscle patterns carrying nearly all task-relevant information for a variety of whole-body reaching movements.

## 1 Introduction

Human motor control has been hypothesized to rely on a modular organization of muscle activity (the so-called muscle synergies or motor primitives) since Bernstein Bernstein (1967) and the seminal works of Bizzi’s group (Bizzi et al, 1991). This hypothesis postulates that the central nervous system (CNS) exploits modularity as a simplifying mechanism to control the many neuromusculoskeletal degrees of freedom underlying goal-directed voluntary movements (Flash and Hochner, 2005; Bizzi et al, 2008). The standard approach to studying motor modularity consists in recording electromyographic (EMG) activity during performance of various motor tasks and then applying dimensionality reduction techniques to decompose the EMG signals into a set of putative synergies. This approach has held substantial evidence supporting the modularity hypothesis (Berniker et al, 2009; Overduin et al, 2012; Nazarpour et al, 2012a; Berger et al, 2013) as well as contradicting (Kutch et al, 2008; Valero-Cuevas et al, 2009) or questioning it (de Rugy et al, 2013; Zelik et al, 2014; Inouye and Valero-Cuevas, 2016).

Arguably, an effective modular decomposition must allow reconstructing the original muscle patterns with good approximation. However, it is known that small errors may be amplified during the control of a nonlinear dynamical system such as the musculoskeletal plant. Therefore, without an accurate model of the body apparatus, complementary indirect analyses are necessary to investigate muscle synergies. In particular, actions are defined in task space and an evaluation of the validity of muscle synergies may conceivably require relating them to task parameters (Nazarpour et al, 2012b; Ting et al, 2012; Todorov et al, 2005). Hence, our approach here is to evaluate modularity models from these two complementary points of view. On the one hand, modularity models are typically assessed based on their ability to approximate the recorded EMG data. However, an absolute expectation on the EMG data reconstruction (quantified as variance accounted for, VAF) is quite arbitrary as high VAF values may also result in overfitting, i.e. the resulting decomposition may contain modules that explain task-irrelevant variance in the EMG recordings, which can be considered as “noise”. In other words, VAF measures focus on approximating EMG activations and ignore the task-relevance of the resulting representations. On the other hand, the modular control theory also implies that the CNS must be able to map a desired motor task onto an adequate activation of modules. For distinct enough motor tasks for which EMG signals differ beyond measurement noise, this suggests that the projection of muscle patterns onto invariant synergies should preserve task discriminability. This inquiry requires contrasting between-task and within-task variability (that may arise from the system’s redundancy and a minimal intervention principle (Todorov and Jordan, 2002)) and assessing the extent to which distinct goal-directed movements are discriminable from the way modules are activated on single movements. Without this property, the relevance of a modular description of muscle patterns would be questionable. Indeed, even though it may yield a very good data approximation, it would mean that the description of muscle patterns in synergy space actually diminished between-task differences in such a way that it becomes hard to decipher which task was really performed on each single trial (Delis et al, 2013b; Alessandro et al, 2013b; Delis et al, 2015).

In this study, we examine a modular description of muscle patterns and compare it with non-modular structures of comparable dimensionality in terms of these two evaluation metrics (VAF and single-trial task decoding). We combine a) a highly comprehensive experiment with b) a unifying modularity model of EMG activity (Delis et al, 2014). Our experimental design comprises surface EMG recordings from a large number of muscles (30) spread across the human body on both hemibodies. Importantly, muscle activity is recorded during performance of a large number of whole-body pointing movements (72 distinct motions or “tasks”) in the 3-dimensional space (Stapley et al, 2000; Leonard et al, 2009). This protocol imposes no further constraints and spans a wide range of movements requiring whole-body coordination including upper and lower limbs while preserving equilibrium. Furthermore, multiple repetitions (30) of the movements are recorded, which allows considering within-task variations and contrasting them with between-task variations in synergy space. In particular, we apply a generic model of modularity, named space-by-time decomposition, which assumes the concurrent existence of spatial and temporal modules (Delis et al, 2014). The use of such a unifying model limits the dependence of conclusions upon the decomposition model used, as temporal, spatial or spatiotemporal modular decompositions have been commonly assumed separately before (Bizzi et al, 2008; Ivanenko et al, 2005; d’Avella et al, 2006). To compute task decoding scores, we employ a single-trial task decoding analysis described in depth in previous studies (Delis et al, 2013b,a) to assess how well we can distinguish the task performed in synergy space, which would be impossible to estimate if trial-averaged data or a limited set of tasks were considered. To compare the VAF and decoding scores obtained with other values obtained from more simple representations of the muscle patterns, we also consider “non-modular” alternative descriptions of the data. The aim of this analysis is to assess whether assuming an advanced model of modularity involving optimization algorithms yields a gain compared to describing muscle activity using parameters directly taken from EMG signals. These analyses show the effectiveness of the space-by-time decomposition model in both approximating muscle patterns and explaining between-task differences in a low dimensional space, during a variety of whole-body pointing movements.

## 2 Materials and Methods

### 2.1 Experimental procedures

#### Subjects

Four healthy participants (2 males and 2 females, aged = 25 ± 3 years old, height = 1.72 ± 0.08 m, weight = 70 ± 7 kg, all values presented hereafter refer to mean ± s.e.m.) voluntarily agreed to participate in this study and performed the experiment. None of them had any previous history of neuromuscular disease. All subjects were made aware of the protocol, and written consents were obtained before the study. Experimental protocol and procedures were approved by the Dijon Regional Ethics Committee and conducted according to the Declaration of Helsinki. As the study focused on intra-individual analyses, few subjects were included in the study.

#### Motor task

Participants were asked to execute whole-body point-to-point movements in various directions at a self-selected pace. The experimental protocol (illustrated in Figure 1) specified 9 targets on 3 vertical bars. Subjects stood barefooted and performed pointing movements between all pairs of targets, termed *tasks* throughout the paper, using the index fingertip of their dominant arm (right) while standing (i.e. a total of 72 different tasks). No constraint was imposed on the left arm. Each participant repeated each task 30 times for a total of 2160 recorded movements per participant. Given the large amount of movements, we separated the whole experiment in two sessions (approximately 2 hours for each session) to avoid participants’ fatigue, with 24h between the two visits. Movements were pseudo-randomized: the same movement was never repeated successively and the same number of trials for each of the 72 motion directions (15 on day 1 and 15 on day 2) were performed in each block. We marked electrode placement on participants’ skin to limit measurement noise due to recording position changes. As reported previously, EMG recordings from different days yield highly similar modular structures (Santuz et al, 2016). Here, we also verified that the removal of electrodes between the two sessions did not critically affect the EMG recordings as well as the identified modules. We computed the correlations between each module of the first session and each module of the second session, separately for temporal and spatial modules. We found a highly significant mean correlation coefficient of 0.89 ± 0.09 for spatial modules and 0.99 ± 0.01 for temporal modules (*for each correlation: p*<0.01) between the two recording sessions of each subject, which shows that the extraction method was robust and that a single extraction including both sessions could be performed. We therefore present the extraction with the 30 repetitions in the Results section.

**Figure 1:**
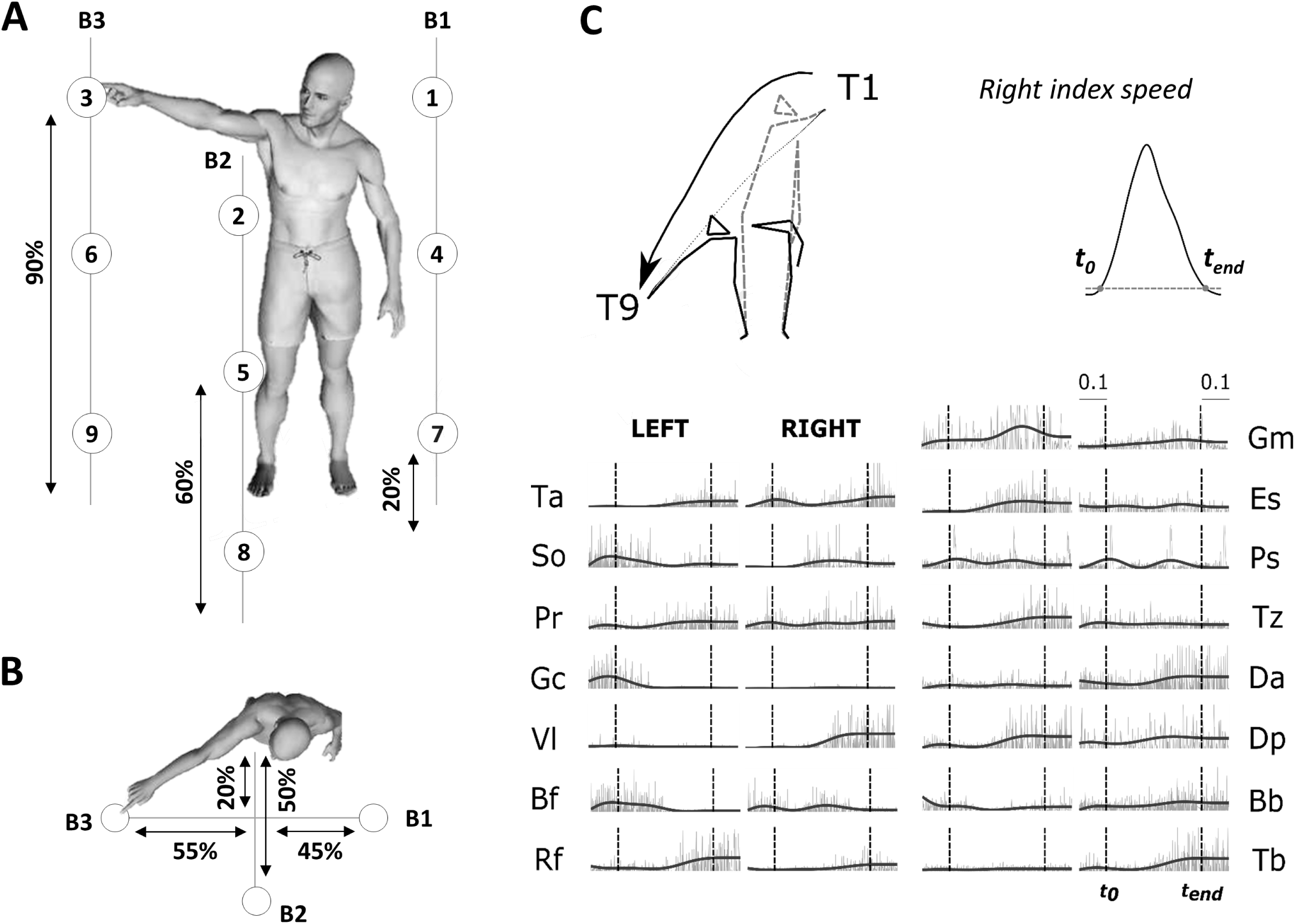
A. Illustration of the experimental protocol. Placement of the 9 targets and the position of the three bars supporting the targets are based on the subject’s height (shown as percentages in figure). Subjects performed point-to-point movements between all pairs of targets (a total of 72 tasks) and repeated each movement 30 times for each direction. B. Top-view of the task. C. Typical raw electromyographic data, for subject S2 and condition T1-T9 (illustrated by the kinematic of initial and final postures and finger trajectories recorded for the associated typical trial). The recorded activity of 30 muscles, normalized in amplitude (divided by the maximum across the whole experiment for each muscle), are plotted from time to-0.1sec to time t_end_+0.1sec. Raw activations are shown in gray and filtered signals, used as input to the module extraction algorithm, are shown in black. Movement onset to and offset t_en_d (chosen as the initial and final time-point of a time period in which the fingertip velocity was continuously above 5% of its maximum) are shown in the figure as dotted vertical lines for each muscle.

#### Kinematics and EMG recording and preprocessing

##### Kinematics

We recorded the 3D positions of 20 retroreflective markers (diameter =
20 mm) using an optoelectronic measuring device (Vicon Motion System, Oxford, UK) at a sampling frequency of 100 Hz. 16 passive markers were fixed symmetrically on the two hemibodies (acromial process, humeral lateral condyle, ulnar styloid process, apex of the index finger, greater trochanter, knee interstitial joint space, external malleolus, and fifth metatarsal head of foot). We added external cantus of the eye on the right face, auditory meatus on the left, and head apex and the first thoraci: vertebra (T1) at the middle. The kinematic data were low-pass filtered (Butterworth filter, cut-off frequency of 20 Hz) and numerically differentiated to compute tangential velocity and acceleration of each marker. To restrict our analysis to movement-related activity, we defined movement onset (*t*_0_) and end (*t_end_*) times as the beginning and end of a time interval in which the fingertip velocity was continuously above 5% of its maximum, and which contained this maximum (Figure 1; Delis et al, 2013b). The average duration of the pointing movements was 1363 ms (±147) with a minimum of 1055 ms (±189; T3->T6) and a maximum of 1635 ms (±305; T9->T1).

##### EMGs

We simultaneously recorded the activations of 30 muscles by means of an Aurion (Milan, Italy) wireless surface electromyographic system. The skin was shaved before electrode placement, and abraded softly. EMG electrodes were placed symmetrically on the two sides of the body on the following muscles: tibialis anterior (Ta), soleus (So), peroneus (Pr), gastrocnemius (Ga), vastus lateralis (Vl), rectus femoris (Rf), biceps femoris (Bf), gluteus maximus (Gm), erector spinae (Es), pectorialis superior (Ps), trapezius (Tz), anterior deltoid (Da), posterior deltoid (Dp), biceps brachii (Bb), triceps brachii (Tb). These muscles were chosen because they are involved in whole-body reaching, and importantly, they not only cover a large part of the human body but they are also easily recordable via a surface-EMG systems. Correct electrode placement was verified by observing the activation of each muscle during specific movements known to involve it (Kendall et al, 2005). During this procedure, EMG signals were monitored in order to optimize recording quality and minimize cross-talk from adjacent muscles during isometric contractions. The trial definition (time interval from *t*_0_ to *t_en_d*) captured the principal EMG signal variations related to the considered conditions. For each trial, the EMGs were rectified, low-pass filtered to obtain smooth envelopes of EMG activity (Butterworth filter, cut-off frequency of 3Hz, zero-phase distortion;Ivanenko et al, 2004) and normalized to 1,000 time steps. A final waveform of 50 time steps was then obtained by using trapezoidal integration of the latter signal on a uniform temporal grid, i.e. we binned the timecourse of the signal into 50 bins and computed the area under the curve in each bin. Movement artifacts were visually removed by discarding the associated trials (<2% of the total number of trials). The data were then normalized in amplitude on a muscle-per-muscle basis by dividing each single-trial muscle signal by its maximal value attained throughout the experiment. A potential detachment of EMG electrodes was assessed, for each subject, by visually checking a posteriori that none of the recorded muscles showed an abnormal change in signal amplitude across trials. For each subject, we finally formed an EMG matrix of (50 time steps × 30 muscles) in rows and 2160 trials in columns consisting of all the movement-related EMG activity (rectified and filtered) of the 30 muscles for all recorded trials. This matrix was used as input to the modular decomposition algorithm to characterize the spatial and temporal structure of muscle activations for this set of movements. Figure 1 (right-down panel) presents movement kinematics (initial and final posture as well as fingertip trajectories) and (both raw and filtered) EMG signals for one pointing condition T1-T9 (diagonal movement from top right to bottom left). Main results were qualitatively the same when defining trials from *t*_0_ − 100 to *t_end_* + 100 to account for the electromechanical delay between EMG activity and real force production.

### 2.2 Space-by-time modular decomposition of muscle activity

#### Space-by-time decomposition model

To represent muscle activity as a structured modular decomposition, we used a space-by-time decomposition model (Delis et al, 2014). This modularity model decomposes muscle activity in separate but concurrent spatial and temporal modules and combines them in single trials using scalar coefficients in order to approximate the recorded EMG activity. More formally, according to the space-by-time decomposition, a single-trial muscle pattern 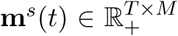 can be written as follows (*s* representing each trial and *T* and *M* being the number of time frames and muscles, respectively):

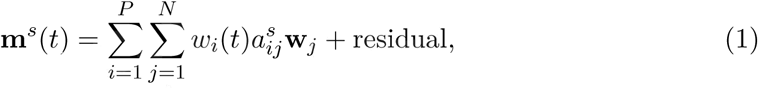

where 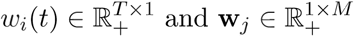 are the temporal and spatial modules respectively, and 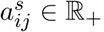 is a single-trial scalar activation coefficient. The free parameters *P* and *N* correspond to the number of temporal and spatial modules, respectively, and are set by the user. The dimensionality of the synergy space is thus *P* × *N*.

#### Variance accounted for (VAF)

To assess how well the space-by-time decomposition approximates the recorded EMG activity, we computed the variance accounted for (VAF) by the space-by-time decomposition. VAF is defined as the total reconstruction error normalized by the total variance of the dataset as follows:

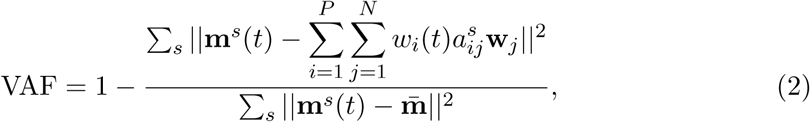

where *S* is the total number of trials, **m̅** is the mean muscle activity across all trials, time steps and muscles and ||.|| denotes the Frobenius norm. Note that different VAFs could be defined by replacing **m̅** by zero or any other reference value (Torres-Oviedo et al, 2006), indicating that VAF values may differ depending on the precise definition of **m̅**.

#### Module extraction

To extract the space-by-time representation of muscle activity in the set of movements under consideration, we applied sNM3F, a Non-negative Matrix Factorization (NMF) based module extraction algorithm that implements effectively the space-by-time decomposition and identify meaningful spatial and temporal modules (Delis et al, 2014). The advantage of NMF-based decompositions is that they restrict the extracted modules and activations to be non-negative, which makes them physiologically relevant for EMG signals reflecting well the “pull only” behavior of muscles (i.e. muscles cannot be activated “negatively”). We input the preprocessed EMG matrix (see above) of each subject to sNM3F and extracted *P* temporal modules, *N* spatial modules and *P* × *N* × *S* activation coefficients capturing all the variations across trials and tasks. The numbers of spatial and temporal modules (*P* and *N* respectively) are free parameters of the algorithm, thus we varied *P* = 1,…, 10 and *N* = 1,…, 10 and computed the decomposition for all the 100 possible (*P*, *N*) pairs. The smallest set of modules describing best the data was then estimated using a decoding analysis (see *Module selection and clustering*).

### 2.3 Task decoding analysis

We complemented VAF evaluation by an additional metric allowing to quantify the plausibility of muscle synergies in task space. Our aim was to assess the reliability of the mapping between module activations and task performance. To quantify this, we used a single-trial decoding analysis that evaluates how well the single-trial coefficients of individual synergies are able to discriminate the 72 different tasks.

The single-trial activation coefficients 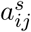 were used to predict the task performed in each trial by means of a linear discriminant analysis (LDA) in conjunction with a leave-one-out cross-validation (Delis et al, 2013b). We quantified decoding performance as the percentage of correctly decoded trials and reported results in the form of confusion matrices. The values on a given row *i* and column *j* of the confusion matrix *C*(*i*, *j*) represent the fraction of trials on which the executed task *j* was decoded as the task *i*. If decoding is perfect, the confusion matrix has entries equal to one along the diagonal and zero everywhere else. The large number of tasks in this study gives a low chance level decoding value (equal to 1/72=1.4%).

### 2.4 Module selection and clustering

#### Selecting the number of modules

Based on the decoding performance evaluation, we used a previously validated automated procedure to select the most compact and task-discriminative space-by-time decompositions (Delis et al, 2013b,a). Our rationale is that VAF includes both the “interesting variance” (related to variations in synergy recruitment across tasks) and the “less interesting variance” (unrelated to variations in synergy recruitment across tasks and that may partly reflect different sources of noise or within-task variability). The presence of the latter variance may make difficult the selection of the correct number of synergies. Also, the removal of “noise” variance using VAF requires the experimenter’s intuition and partly arbitrary or ad-hoc criteria (Delis et al, 2013b,a). This complementary evaluation overcomes this problem by singling out (by single-trial task decoding) only the task-discriminating variance and then studying the dependence upon the number of synergies of this part of the variance.

We thus implemented the selection of the number of modules as follows. For each subject, we progressively evaluated the statistical significance of the task-discriminating information added when increasing gradually the number of temporal and spatial modules (*P*, *N* respectively) in the decomposition model. The number of modules was then selected as the step at which adding a supplementary module did not give any significant decoding gain (*p* > 0.05). To assess the significance of decoding performance, we employed a permutation test where we randomly shuffled the coefficients corresponding to the added module (while the distributions of all other coefficients were unaffected) and computed discrimination performance. For instance, for a given value N, we compared the decoding performance of the synergy parameters when using the N synergies with the decoding performance of the parameters of all subsets consisting of N—1 synergies plus the parameters of the N-th synergy pseudo-randomly permuted (“shuffled”) across conditions. We repeated this shuffling procedure 100 times to obtain a non-parametric distribution of decoding performance values in the null hypothesis that the additional synergy does not add to the decoding power of the synergy decomposition. This procedure ensured the detection of modules that explain the “task-relevant” variability and the exclusion of other sources of noise (“task-irrelevant” variability) (for a more detailed description, see Delis et al 2013b,a).

#### Clustering analysis

To compare modules of the same type (spatial or temporal) extracted from different subjects, we grouped them using an agglomerative hierarchical cluster analysis (Hastie et al, 2009). Although it was not crucial for the present study, such a clustering can be useful for visualization purpose and for comparison of our results with other studies. In particular, it is worth mentioning that the modular control hypothesis does not impose that different subjects must have the same modules but that states that each subject may rely on its a modular structure to generate genuine muscle patterns. In the following, we will present the procedure used for clustering in detail for spatial modules, but the same procedure was followed also for clustering the temporal modules. We quantified the similarity between spatial modules *i* and *j* as their correlation coefficient (*R_i,j_*). We considered spatial modules as *M*-dimensional vectors and computed correlation coefficients between all pairs of modules across all pairs of subjects. Using *R_i,j_* as distance measure, we created a hierarchical cluster tree from all module pairs (Matlab function “linkage” with the “average” distance method, i.e., using as distance between two clusters the average distance between all pairs of objects across the two clusters). The number of clusters was set to the maximum number of spatial modules across subjects (i.e. 7 here). The correlation between modules was then computed as the mean pairwise correlation between all pairs of modules within each cluster.

### 2.5 Signifiance of identified decompositions

We used a permutation test to assess the ability of the identified decompositions to uncover meaningful structure in the data. We compared the VAF and decoding performance of the identified decompositions with the VAF and decoding performance values obtained when decomposing structureless data. We generated structureless data from the recorded data by randomly permuting the muscles for each time step of each trial in every movement. The input matrix thus had exactly the same numerical values but was devoid of biomechanical significance. For each subject, we performed 10 different permutations, which resulted in 10 simulated datasets on which we applied sNM3F to extract space-by-time decompositions and computed VAF and decoding performance of the resulting decompositions. We considered as significance level the maximum of the VAF and decoding performance obtained for these decompositions. Quality of the VAF and decoding performance obtained for the recorded data was then evaluated relative to this significance level.

### 2.6 Comparison with non-modular muscle pattern descriptions

To compare the efficiency of the extracted modular decompositions with non-modular alternatives, we computed decoding performance and VAF of non-modular descriptions of the data with equal number of parameters as the modular decompositions. In particular, we examined whether alternative descriptions of muscle activity that do not rely on an explicit modularity model are more or less effective than the space-by-time decomposition in a) discriminating the task performed and b) approximating the EMG signals. This analysis also served to investigate whether a subset of the recorded muscles or a shorter temporal window of muscle activity suffices for the characterization of a) the recorded EMG signals and b) the task differences in single-trials. Thus, we compared task decoding and VAF results of the space-by-time decomposition with those obtained by artificially reducing the spatial dimensions (i.e. choosing a subset of muscles) or the temporal dimensions (i.e. splitting muscle activity into shorter temporal windows) or a combination of the two. To this aim, we divided the muscle activity of each of the *M* muscles (spatial dimension) into *B* bins (temporal dimension) and computed the root-mean-square (rms) of the EMG signal of each muscle within each temporal bin. This procedure yielded *M* × *B* parameters for each trial, which we refer to as “non-modular” parameters. This gives an approximation of the EMG signals that does not rely on any modularity model that can be straightforwardly compared to the approximation achieved by the modular model. In fact, the non-modular approximation relies on parameters extracted directly from the signals and when the number of parameters reaches the number of data points, it becomes a perfect reconstruction of the original signal. Hence, by gradually reducing the number of parameters, we can obtain a fair comparison between the modular model performance and the non-modular approximation. To perform a fair comparison, we then matched the number of non-modular parameters with the dimensions of the space-by-time decomposition, i.e. selected an identical number of parameters for both descriptions, in three different ways.

Firstly, we examined whether adding more spatial dimensions (and ignoring the temporal structure of the data) would enhance task discrimination performance. Thus, the first set of non-modular parameters described only the spatial dimension of muscle activity by varying the number of muscles retained (*M* = *N* × *P*) and keeping only one temporal dimension (*B* = 1). Secondly, we examined the effect of adding more temporal dimensions (and ignoring the spatial structure of the data). Thus, to obtain the second set of non-modular parameters, we varied the number of temporal bins (*B* = *N* × *P*) and kept only one spatial dimension (*M* = 1). Thirdly, we selected equal numbers of spatial and temporal dimensions with the space-by-time decomposition (*M* = *N*, *B* = *P*) and asked whether we could achieve higher task discrimination using the non-modular parameters instead of the modular parameters. We repeated parameter selection for each of the three sets 20 times (by randomly selecting muscles and/or bins, when appropriate). We used these three sets of parameters to compute the decoding performance of the non-modular muscle activity descriptions and compare it with the decoding performance of the space-by-time decomposition. Note that the VAF could not be evaluated from some these non-modular decompositions because they involve only a subset of the recorded muscles; thus, they can reconstruct only a part of the EMG recordings.

To resolve this issue, we then quantified the maximal decoding and maximal VAF that can be achieved by the dataset under investigation using non-modular descriptions of the data involving all recorded muscles. We described EMG activity in single trials using the rms values from the single-trial recordings of all *M* = 30 muscles, binned into increasing numbers of temporal bins (*B* varying from 1 to 50). We input these values to LDA and computed the maximal “non-modular” decoding performance (for *B* = 50 the non-modular description was identical to the one given by the original data set). Computing the VAF was also possible here, using the following procedure: the reconstructed EMG matrix of this non-modular decomposition was obtained by assuming all time points within each bin to be equal to the rms value of that bin (i.e. a piece-wise constant function). The resulting reconstructed data matrix 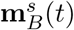 for each trial had equal dimensions as the original single-trial EMG data matrix **m**^*s*^(*t*) and was defined as follows:

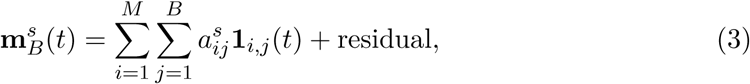

where **1**_i,j_(*t*) € ℝ^*M*^ is the indicator vector function that is equal to 1 on the *i^th^* component if *t* belongs to bin *j* and to 0 elsewhere, and 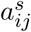 is the rms value of muscle *i* for bin *j* and trial *s*. Hence, the VAF of the non-modular decomposition can be computed from Eq. 2 by replacing the double sum term by 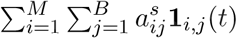.

**It is worth mentioning that the non-modular models tested here may be sub-optimal in terms of VAF or decoding performance. Indeed, for the same number of parameters, better results could be obtained by selecting different spatiotemporal representations of the muscle patterns. The subsequent comparison will be done between the results achieved by the modular model, which maximizes VAF and results achieved by the non-modular models that do not attempt to maximize neither the VAF nor the decoding scores. The choice of the non-modular descriptions used in the present study was guided by simplicity and to provide a basis of comparison.**

## 3 Results

### 3.1 Basic kinematic features

The four subjects performed 72 different point-to-point movements between pairs of targets among 9 predefined target points. All finger velocity profiles appeared to be bell-xsxsxshaped across subjects and conditions, as previously observed in whole-body reaching movement (Thomas et al, 2005; Berret et al, 2009). Targets were attained with an overall mean spatial error of 10mm ±2mm (from 8 to 15 mm for T6-T3 and T9-T5 respectively). Raw EMG data, associated with the task T1-T9, for a typical trial of subject S2, are shown in the Figure 1. These recordings for all conditions and trials formed the EMG matrix that was used for module decomposition.

### 3.2 Low-dimensional modular decomposition in space and time

We extracted a space-by-time representation of muscle activity by applying the sNM3F algorithm to the EMG recordings of each subject. Figure 2 illustrates the VAF and decoding performance graphs (upper surfaces in all plots) as a function of the number of spatial and temporal modules for one typical subject. These graphs provide insights about the number of spatial and temporal dimensions that are necessary to describe the set of tasks at hand. For all subjects, VAF exhibits a smooth increase with the number of temporal and spatial modules with no clear saturation point. In contrast, task decoding performance grows quickly and reaches a plateau for all subjects. The 3D decoding graphs show a larger effect of the spatial dimension on decoding compared to the temporal one indicating that precise muscle groupings may carry more task-related information than precise timing of muscle activations. The sets of temporal and spatial modules were selected as the dimensions of the space-by-time decompositions for which no statistically significant gain (*p* < 0.05) in decoding was obtained when adding more (spatial or temporal) modules. In particular, four temporal modules (*P* = 4) appear to carry all decoding power for all subjects, whereas the number of spatial modules varies across subjects and is usually higher (S1: *N* = 4, S2: *N* = 6, S3: *N* = 7, S4: *N* = 5). The resulting decompositions achieved on average across subjects (mean ± sem) a VAF value of 68% ±5% and decoding performance of 86% ±1%. The corresponding graphs for all subjects are presented in Figure 2. VAF values may appear relatively small compared to other studies, especially if one sets a somewhat arbitrary threshold such as 90% for selecting the number of modules (Torres-Oviedo et al, 2006; Hart and Giszter, 2004; Ting and Macpherson, 2005). This discrepancy is partly due to the fact that data were not averaged across trials here (for comparison, the VAF obtained from averaged data was 90% ±6% when considering the optimal number of modules for each subject, see Discussion for more details on these VAF differences). We assessed the statistical significance of these VAF values by performing a permutation test (see Methods for details on this computation). The lower surfaces in each plot of Figure 2 represent significance levels for VAF and decoding values for unstructured data, which we compared to the ones obtained from the space-by-time decompositions. For the selected number of modules, significance level for VAF is 9% ±3% and for decoding performance 19% ±5% across subjects. Note that, for decoding, significance level is higher than theoretical chance level (˜1.4%) because our permutation technique preserved the order of trials and tasks (only muscles were shuffled for each time step). Overall, VAF and decoding scores were significantly larger than their corresponding chance and significance levels. These results validate that the identified space-by-time decompositions account for relevant features of the recorded EMG data and are not just an artifactual output of the methods.

**Figure 2:**
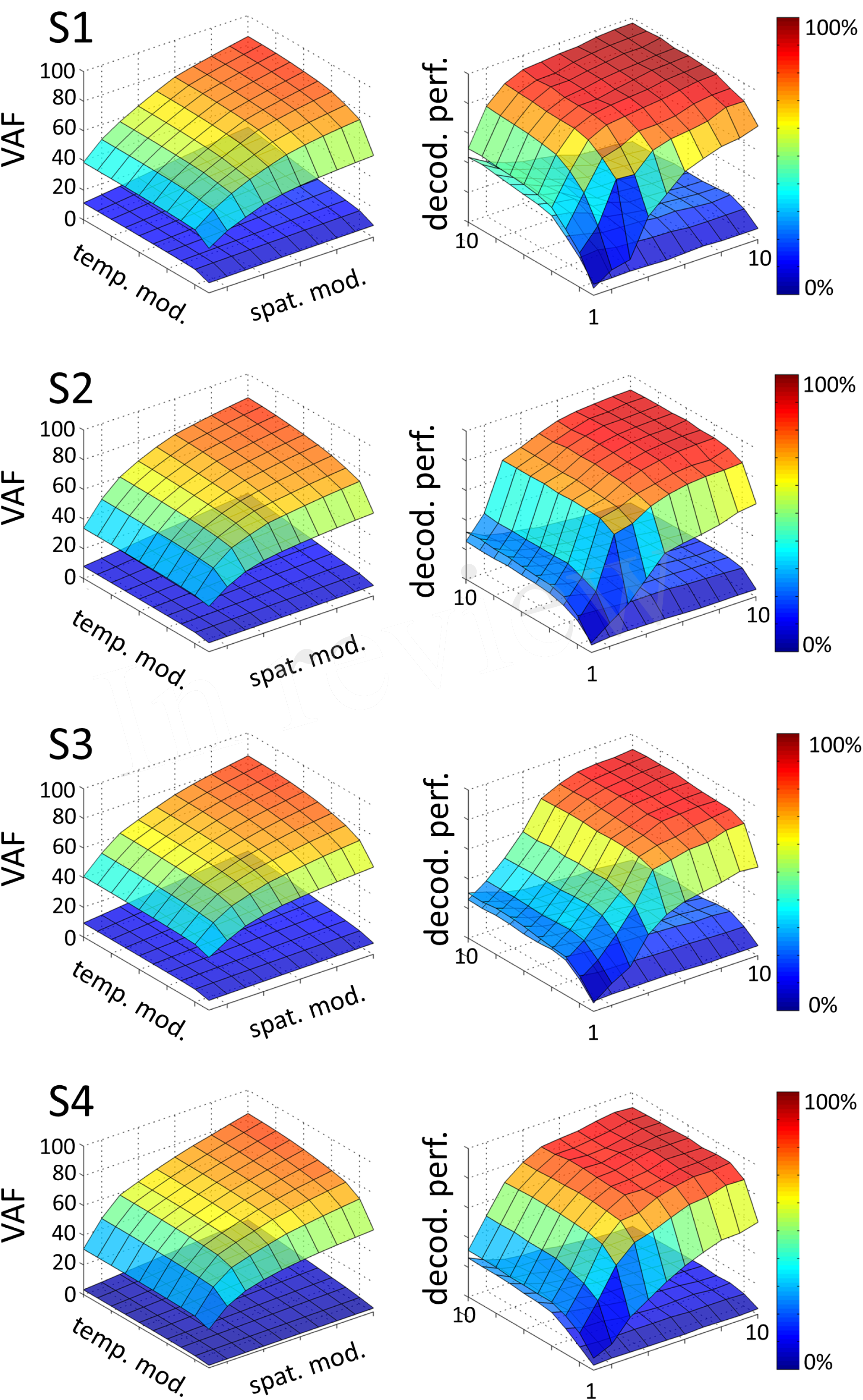
Influence of the number of temporal (*P* = 1… 10) and spatial modules (*N* = 1… 10) on VAF (left) and decoding performance (right), for each subject (S1, S2, S3, S4). In each graph, the upper surface corresponds to the decoding performance and VAF of the space-by-time decomposition as a function of the number of spatial and temporal modules. The lower surface represents significance levels for VAF and decoding values, computing as the maximum decoding and VAF values obtained from a permutation test where synergies were extracted from a random shuffling of the EMG data matrix across muscles.

### 3.3 Consistency, task-independence and generalizability of spatial/temporal modules

We then examined the composition and shape of the extracted spatial and temporal modules, and their similarity across subjects. In the space-by-time decomposition, temporal modules are *T*-dimensional vectors containing time-varying patterns, accounting for the timing of muscle activity. Here, the identified temporal modules were highly consistent across subjects (mean correlation coefficient r=0.92±0.06 in each cluster). Each temporal module was composed of a single activation burst (Fig. 3) and the four modules were successive in time to describe the temporal profile of muscle activity in different temporal windows of the full movement duration, which is a common finding in literature (Ivanenko et al, 2005, 2004; Chiovetto et al, 2010, 2013). This result may be the consequence of the common organization of all movements consisting in an acceleration phase followed by a deceleration phase.

**Figure 3:**
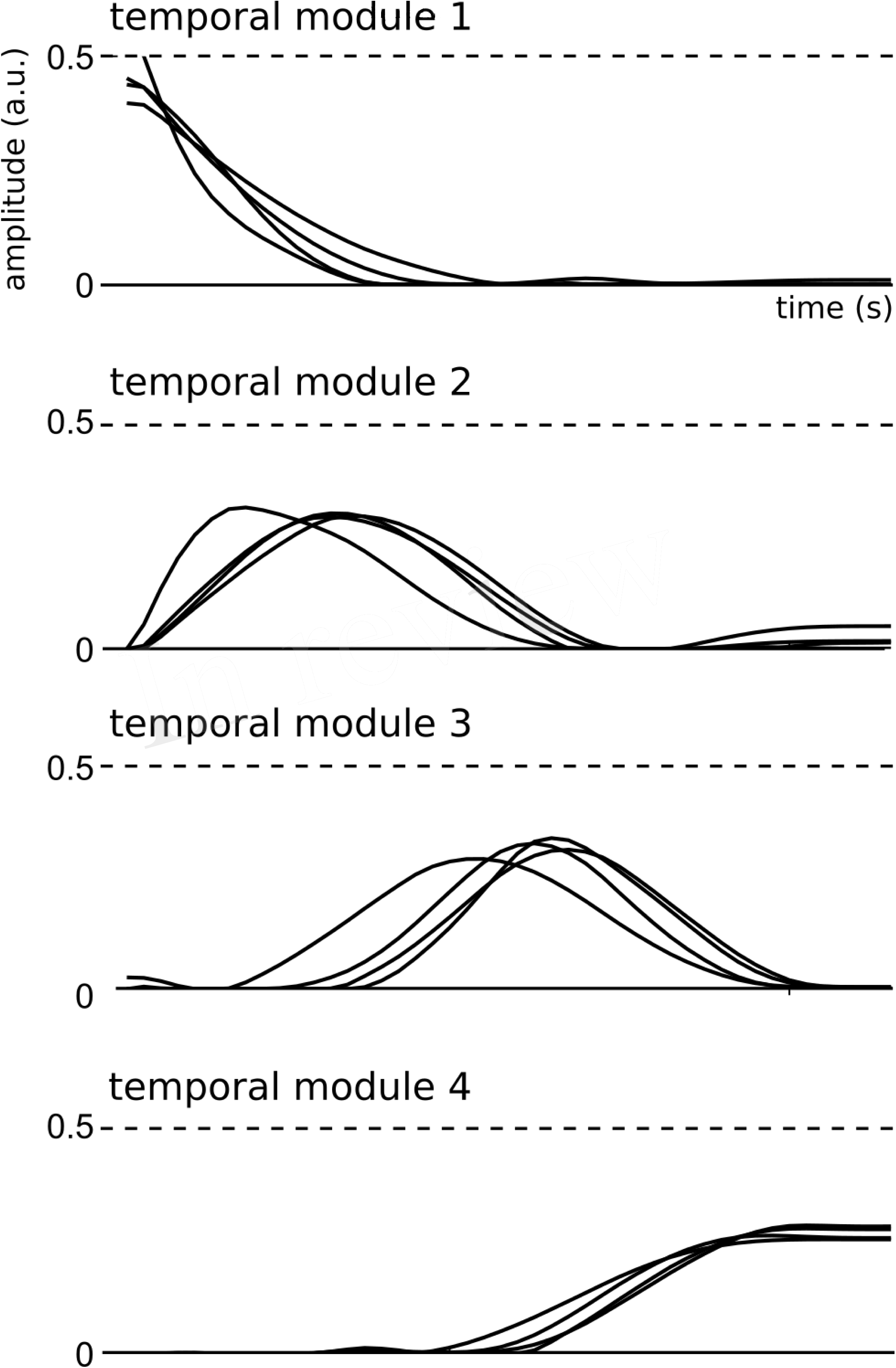
Representation of the four temporal modules identified by the space-by-time decom-position for all subjects.

Spatial modules are *M*-dimensional vectors of muscle activation levels. The identified spatial modules exhibited higher variability across subjects (mean correlation coefficient r=0.92±0.15) than the temporal ones, which might be partly explained as a result of differences in movement kinematics, the subject-specific number of spatial modules (thus requiring different muscle groupings) and/or physical discrepancies between participants (muscle sizes, skin conductance etc.). We determined four clusters of spatial modules across subjects. Each spatial module activated muscles spread across the whole body (and on both hemibodies, Fig. 4). This suggests that the extracted spatial modules represent functional muscle couplings for performing the movement at hand rather than purely anatomical groupings of muscles controlling the same joints or body parts.

**Figure 4:**
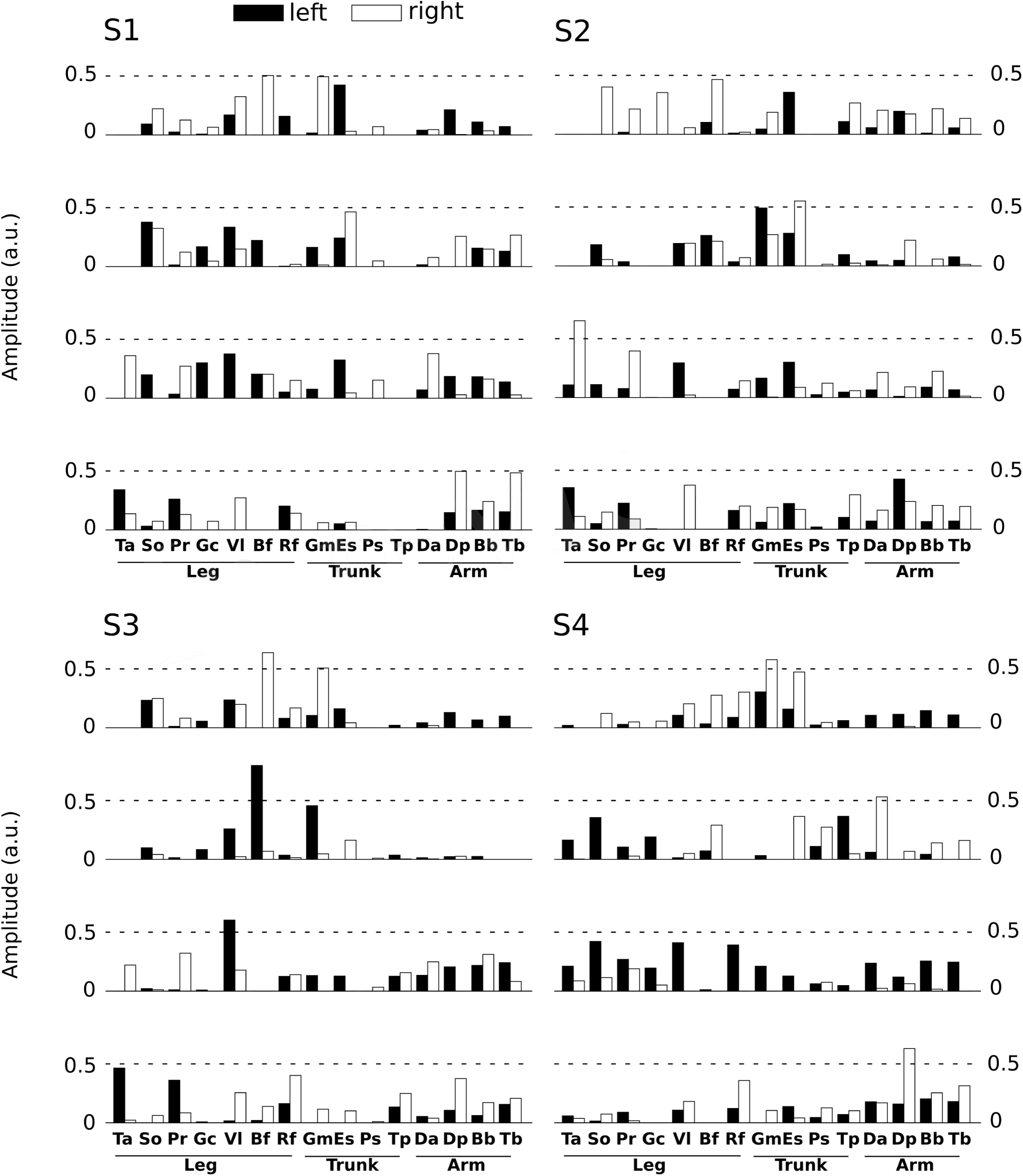
Representation of the main four spatial modules for all subjects. Each panel presents the four first spatial modules of each subject (S1 left-high, S2 right-high, S3 left-low, S4 right-low). Spatial modules were sorted for each subject relative to their similarity with the four modules of S1. Note however that we found more than 4 spatial modules for some subjects, but we only depict 4 of them here for convenience. The 30 muscles are represented by vertical bars (white filled if placed on the right hemibody, black on the left hemibody). For S1 only 27 muscles were recorded, the three absent muscles have zero values in the S1 panel.

To quantify whether the extracted modules were able to generate muscle patterns underlying new tasks, we ran two additional tests. We extracted spatial and temporal modules using only a subset of the motor tasks (75% of all tasks) and then quantified the similarity of these modules with those extracted from the full dataset. To simplify these tests, we ran the decomposition on averaged data (trial-averaged for each task) for each subject and for the optimal number of modules. We first performed the decomposition on a task subset selected randomly to encompass 75% of the total number of tasks (i.e. 54 tasks out of 72). We repeated this process 10 times for each subject. The results showed a high correlation between the modules extracted from subsets of tasks and the initial decomposition (containing all tasks): averaged r values across best-matching pairs of modules and subjects was 0.96±0.03 for the spatial modules, and 0.99±0.01 for the temporal modules. Accordingly, the modular decompositions of the task subsets approximated the test EMG data equally well as the original decomposition that included all tasks (VAF=90% ±6%).

In our second test, we chose specific subsets of tasks to assess the effect of target location on the extracted modules. For each subject, we tested four different subsets (of 54 tasks out of 72): (1) discarding all tasks ending at any target of the top row: r=0.90±0.11 (mean r across pairs of modules and subjects) (2) discarding all tasks ending at any target of the bottom row: r=0.93±0.10 (3) discarding all tasks ending at any target of the left bar: r=0.92±0.07 (4) discarding all tasks ending at any target of the right bar: r=0.98±0.01. Also for this test, the obtained VAF values were on average the same as the original ones (VAF=90% ±6%).

Therefore, the high similarity and good generalization we found in all cases supports the relevance of the extracted modules to approximate and construct genuine muscle patterns.

### 3.4 Efficiency of the identified space-by-time decomposition in task discrimination

In this part, we aimed to assess the effectiveness of the identified space-by-time decompositions with respect to task decoding performance. To this end, we used the single-trial parameters of the decompositions, i.e. the *N* × *P* activation coefficients, to decode which of the 72 tasks was performed on each trial. Decoding results are shown as confusion matrix for all subjects (Fig. 5). Each entry of the confusion matrix *C*(*i,j*) represents the percentage of times task *j* was decoded as task *i*. In Figure 5, only the matrix diagonal shows high values (on average higher than 90%), which indicates highly accurate direction decoding from the way modules are recruited on single trials. We also observe two light blue lines parallel to the diagonal (one above and one below the diagonal) indicating incorrect classifications for some pairs of tasks (corresponding to 11% of decoding errors on average). These decoding errors concerned tasks starting from neighboring points on the same bar and ending at the same point on a different bar, for which the between-task and within-task variability of muscle patterns was likely less distinguishable. In particular, starting points T1, T2 and T3 (higher level) were confused as T4, T5, and T6 (middle level) respectively and vice-versa (see Fig. 1 for target positions). Hence, these confusions suggest that decoding is harder between tasks that have the same spatial direction (left or right) and the same endpoint and their starting locations differ only in the height dimension. Starting points at the lower level were confused less often (<10% decoding errors) probably because of the higher involvement of lower body muscles required for these movements, which distinguishes them from the middle and higher level starting points. To gain more intuition on how activation coefficients are modulated by the task, we also plotted the trial-averaged activation coefficients for each initial and final target (see Figure 6 for a subject S2 and supplementary material A for other subjects). For all subjects, some coefficients are clearly modulated by changes in the final position (e.g. S1:A44, S2:A34, S3:A14, S4:A44), and others by changes in the initial position (e.g. S1:A31, S2:A11, S3:A21, S4:A31). We also observed a few coefficients that were less clearly linked to the modulation of just initial or final position. These coefficients may encode other task parameters. However, verifying this point requires a more thorough decoding analysis on other task parameters (e.g. amplitude, direction) and is out of the scope of the present study.

**Figure 5:**
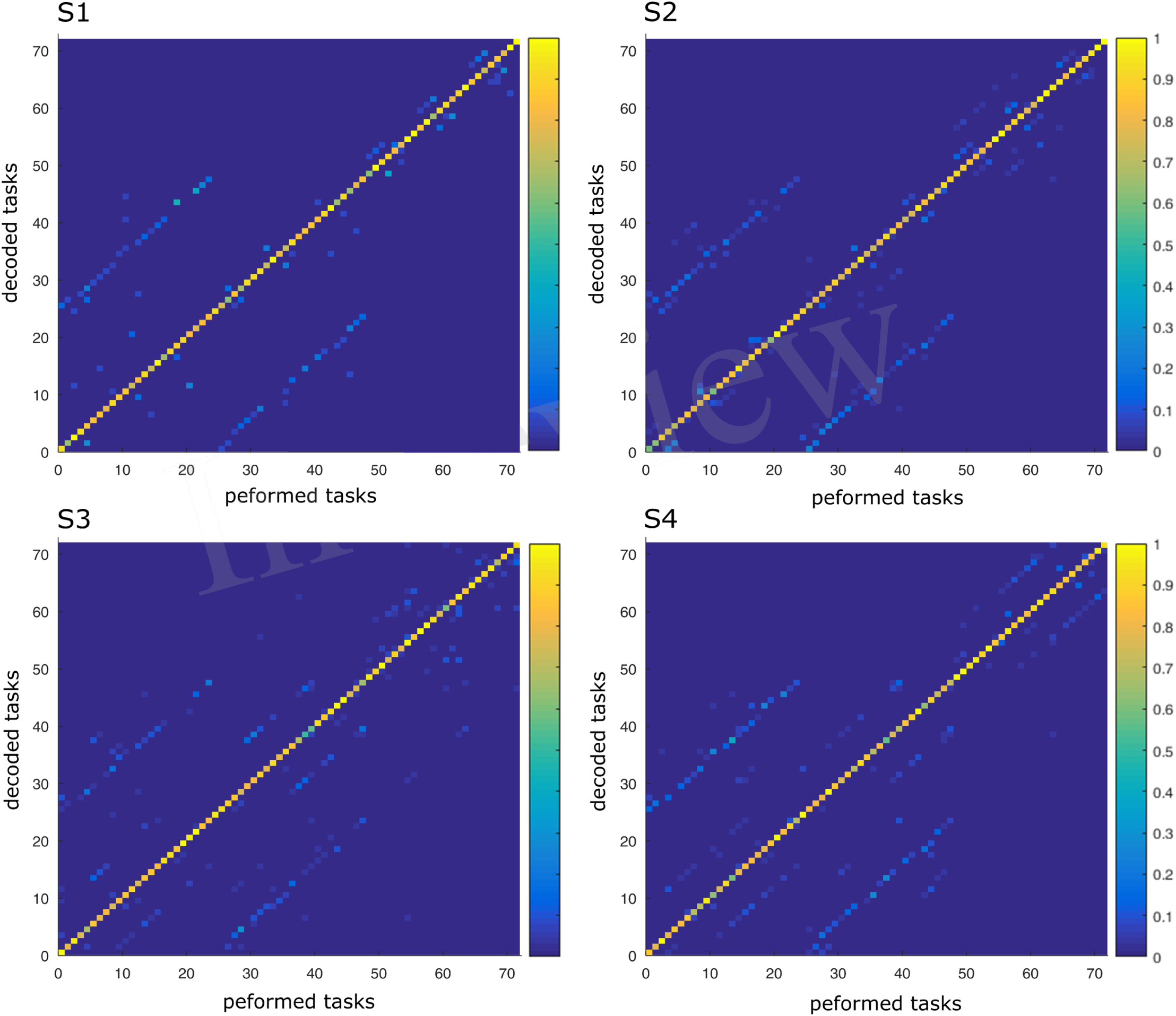
Confusion matrix of the four subjects. For each graph, rows correspond to the decoded task and columns to the task actually performed by the subject. Each color-scaled entry of the matrix *C*(*i,j*) represents the percentage of times the task *j* was decoded as the task *i* (yellow corresponds to 100% correct decoding and blue is 0%).

**Figure 6:**
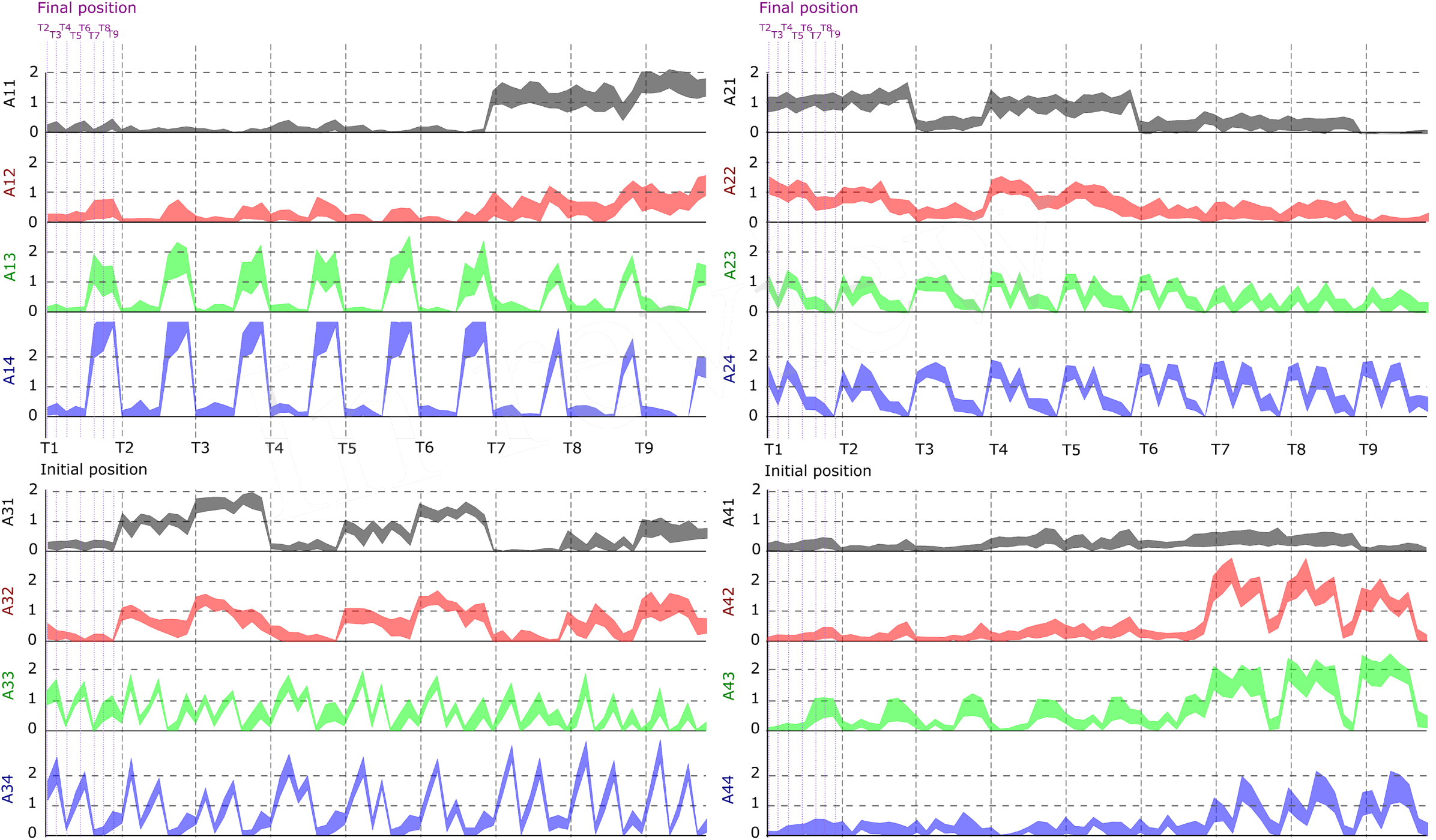
Dependence of the activation coefficients on the initial and final posture. Each graph represents the value of an *a_ij_* coefficient as a function of the initial position (from T1 to T9; grey-dotted vertical bars) and the final position (from T1 to T9; violet-dotted vertical bars), with i being the number associated to temporal modules (from 1 to 4) and j to spatial modules (from 1 to 4). Each panel represents one temporal module (1-left up, 2-right up, 3-left down, 4-right down), each color one spatial module (1-grey, 2-red, 3-green, 4-blue). Results are shown for the data of subject S2.

### 3.5 Effectiveness of space-by-time modularity as compared to non-modularity

To compare the effectiveness of the modular decomposition in terms of task discrimination, we confronted its decoding power with the ones obtained with the same number of parameters but taken directly from the recorded muscle activity, that is, without assuming any advanced modularity model (see Materials and Methods for details on the extraction of non-modular parameters).

Here to treat all subjects with the same methodology, we first computed the decoding performance of the space-by-time decomposition with 9, 16 and 25 single-trial coefficients (i.e. (*N, P*) = (3, 3), (*N*, *P*) = (4,4) and (*N, P*) = (5, 5), respectively, Fig. 7, black curve). We then compared the decoding performance of the modular decomposition with the 95% confidence intervals of decoding performance obtained using three simple sets of parameters of equal dimensionality (red, green and blue areas) that capture the spatial, temporal and spatiotemporal structure of the EMG data respectively (see Methods for details on these computations). For all subjects, the space-by-time decomposition carried higher task discrimination power than the three sets of parameters. Notably, the worst decoding performance was obtained when decoding was based only on the temporal parameters of an individual muscle (red area) suggesting that temporal dimension carries less information about task differences, possibly because all reaches were made of one acceleration and one deceleration phases with similar timings. On the contrary, when exploiting the spatial dimension (overall activation of a set of muscles, green area) decoding results were higher. This finding suggests that direction information was mainly carried by the relative activation levels of different muscles (i.e. the spatial dimension), whereas the precise timing of muscle activations did not contribute to the discrimination of different movements. It is noteworthy that not only was decoding lower for the tested sets of non-modular parameters but also these parameters can reconstruct only a subset of the EMG data and thus VAF cannot be evaluated for them (i.e. they are not able to construct complete muscle patterns).

**Figure 7:**
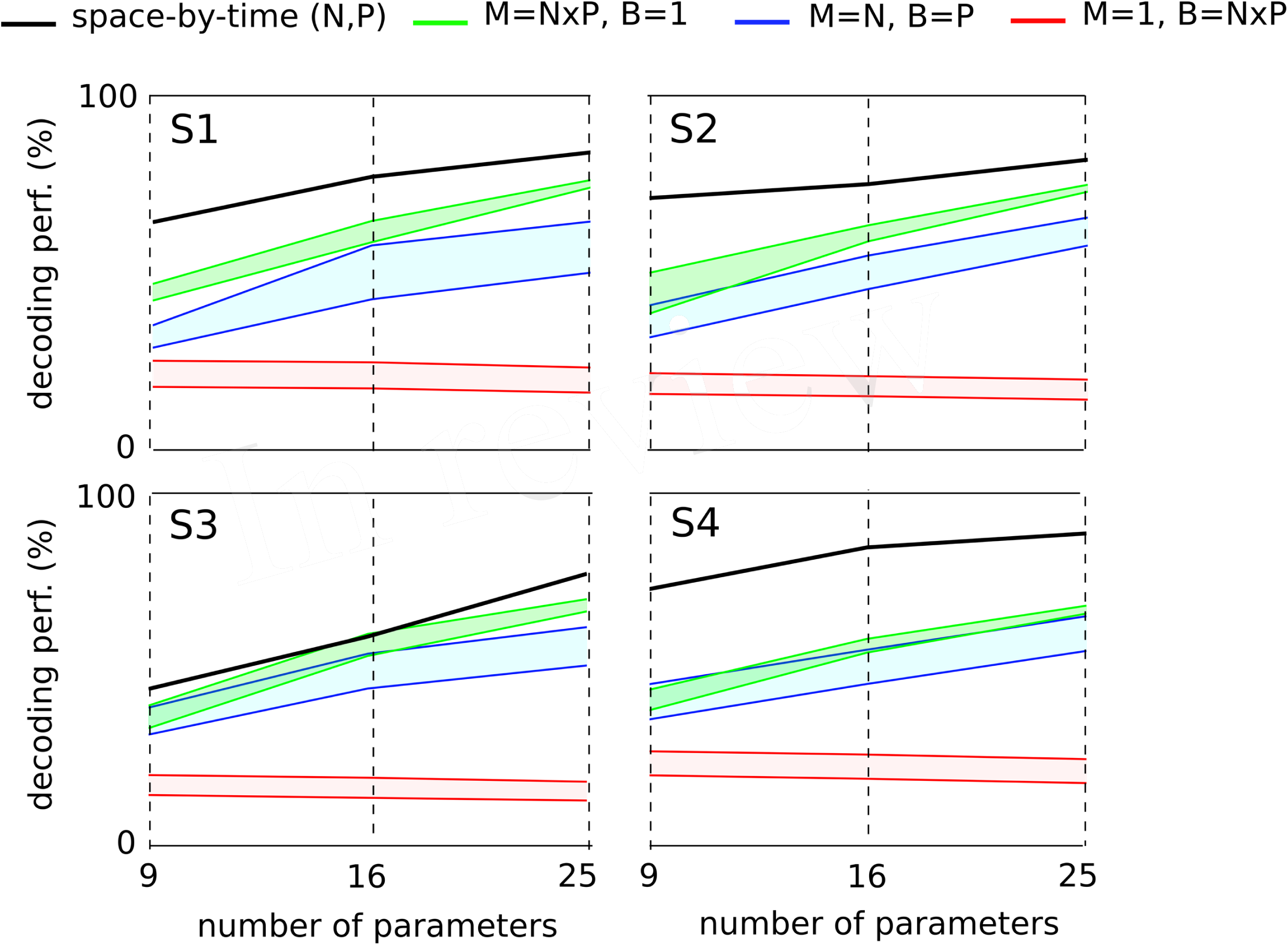
Comparison of decoding power between the space-by-time decomposition and three other sets of EMG parameters of equal dimensionality for each subject. The different sets of parameters were compared in terms of their decoding power for three different dimensionalities (9, 16 and 25 parameters corresponding to (*P*, *N*) = (3, 3),(4, 4),(5, 5) spatial and temporal modules respectively). For each subject’s graph, the black line represents the decoding power of the space-by-time decomposition. For the three other sets of decoding parameters, we plot the 95% confidence interval for the decoding performance results obtained across 20 repetitions. The red area represents the decoding performance for one randomly selected muscle and 9, 16 and 25 temporal bins (*M* = 1, *B* = *N* × *P*). The green area represents the decoding performance for 9, 16 and 25 randomly chosen muscles and one temporal bin per muscle (*M* = *N* × *P*, *B* = 1). The blue area represents the decoding performance for *N* randomly chosen muscles and *P* bins per muscle (*M* = *N*, *B* = *P*). *M* refers to the number of muscles, *B* to the number of temporal bins per muscle and *P*, *N* represent the number of temporal and spatial modules respectively.

In order to be able to also evaluate the VAF, we considered a non-modular model describing muscle patterns with piece-wise constant functions (see Eq. 3). This model included all 30 muscles -as the spatial dimension carries most of the task information- and contained a varying number of bins to test different temporal resolutions. When all bins and all muscles were considered, the model yielded a complete and exact description of the original muscle patterns. We observed that the maximal decoding performance that could be attained with the activation parameters of such a model was 95% ±4% across subjects, with 5 temporal bins (i.e. 150 decoding parameters, black curve in Fig. 8A). Increasing further the number of bins turned to decrease the decoding performance first slightly and then drastically, confirming our previous findings about the small contribution of temporal precision to decoding. In comparison, the maximum decoding performance obtained by the space-by-time decomposition was 91% ±3% (for N=10, P=10, i.e. 100 parameters, gray curve in Fig. 8A). Interestingly, a smaller set of parameters for the modular model (25 parameters, *N* = 5, *P* = 5) already achieved decoding performance of 84% ±5%. For a comparable number of parameters, decoding performance was 86% ±3% for 36 modular parameters (see gray curve in Fig. 8A) versus 75% ±5% for 30 non-modular parameters (see black curve in Fig. 8A). Note that in these analyses only the number of parameters that must be specified for each single movement to create a full muscle pattern are counted. Notably, the extracted space-by-time decomposition using the optimal number of parameters for each subject (S1: *N* = 4, *P* = 4, S2: *N* = 6, *P* = 4, S3: *N* = 7, *P* = 4, S4: *N* = 5, *P* = 4, i.e. 22±1 parameters across subjects) yielded 86% ±1% decoding, whereas the decoding performance of the non-modular decomposition with 30 parameters was 75% ±1%.

**Figure 8:**
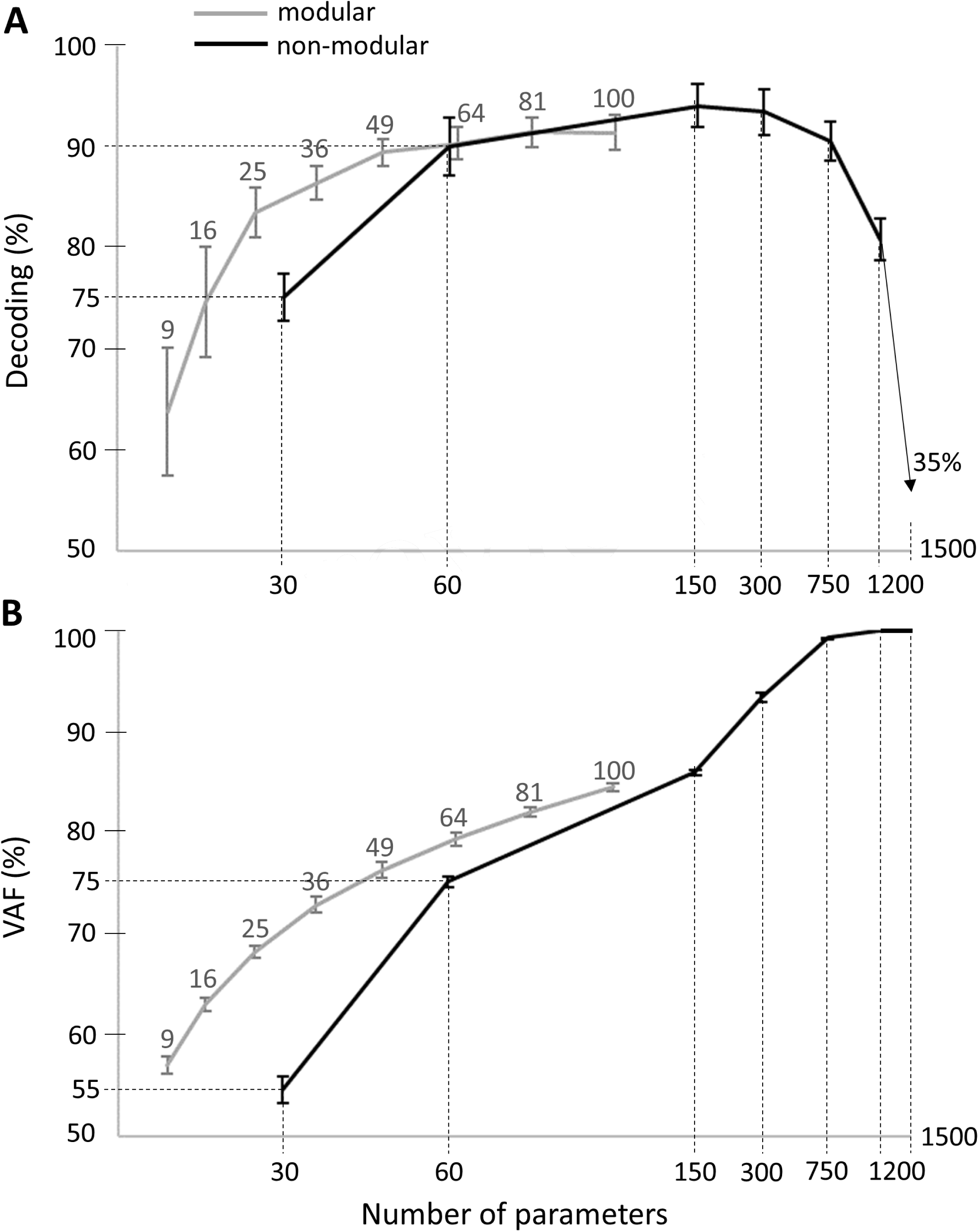
Comparison of decoding performance (A) and VAF (B) as a function of the number of single-trial parameters between the modular and non-modular decompositions. Gray lines represent the average decoding performance and VAF across subjects of the modular decomposition using 3, 4, 5, 6, 7, 8, 9, and 10 temporal and spatial modules, which correspond to a total of 9, 16, 25, 36, 49, 64, 81 and 100 single-trial parameters respectively (top gray x-axis). Black lines represent the average decoding performance and VAF across subjects of the non-modular decomposition. As single-trial parameters of the non-modular decomposition, we used the rms values of the EMG signals of all (30) muscles binned in 1, 2, 5, 10, 25, 50 temporal windows, which corresponds to a total of 30, 60, 150, 300, 750, 1500 single-trial parameters respectively (bottom x-axis).

Regarding data approximation, we observed that increasing the number of bins led to a gradual increase of the VAF values up to a maximum of 100% when the full muscle patterns (50 time bins for 30 muscles, i.e. 1500 parameters specified for each single movement) was used (see black curve in Fig. 8B). For the modular decomposition (gray curve in Fig. 8B) and small numbers of parameters (from 1 to 100), VAF values were qualitatively equal or higher than the ones associated with the non-modular descriptions of muscle patterns (e.g. for 64 modular parameters: VAF = 79% ±1% vs for 60 non-modular parameters: VAF = 75% ±1%). Again, the extracted space-by-time decomposition using the optimal number of parameters for each subject (22 ±1 parameters across subjects) yielded 68% ±1% VAF, which is higher than the VAF of the non-modular decomposition with 30 parameters (55% ±3%). When increasing the number of parameters, the non-modular decomposition achieves higher VAF values reaching 100%, as expected when all data points are used, which is also the case when choosing *N* = *M* and *P* = *T* for the extraction of modules (although it would be meaningless regarding the modularity hypothesis as there would be no muscle grouping and no dimensionality reduction).

## 4 Discussion

In this study, we analyzed the effectiveness of space-by-time modularity in describing muscle patterns underlying a set of complex goal-directed movements. We designed a comprehensive experiment comprising 72 distinct whole-body pointing movements and collected 30 EMG channels. We assessed the extent to which a space-by-time modular decomposition of muscle activity could a) approximate the rectified/filtered EMG data on single trials and b) convey task-relevant information. Although descriptive, the present study shows that space-by-time modularity provides a relevant overview of how muscle patterns may be formed during whole-body voluntary movements, as discussed below.

### 4.1 Parsimonious representation of EMG data in space and time

Four bursts of muscle activation characterized the timing of movement-related EMG activity. Notably, this set of temporal modules was very consistent across subjects and more temporal precision did not improve the characterization of the task performed using decoding analysis. Refining the number of temporal modules only contributed to increasing the VAF, i.e. yielded a better reconstruction of EMG patterns. This finding shows that the task-relevant information is mainly conveyed in four successive temporal recruitments that may correspond to different phases of the goal-directed movement. The nature of the task may have defined the four temporal phases observed here: postural stabilization over starting point, movement initiation, movement deceleration and stabilization over endpoint. The obtained temporal modules have a particular form similar to bursts of muscle activity. It is, however, important to note that no mathematical constraint was imposed on the selection of these modules. In other words, the modules were only constrained to be non-negative and no specific shape was imposed on them (e.g. Gaussian). In addition, similar temporal modules have been described in different studies (different motor tasks and different animal models) using different extraction techniques Ivanenko et al (2004, 2005); Dominici et al (2011); Hart and Giszter (2004), which suggests that the obtained shapes are likely not a by-product of the algorithm or the task constraints but are physiological. This supports the robustness of the identified temporal structure of motor modules irrespective of the study-specific signal preprocessing procedure (Hart and Giszter, 2010; Kargo and Giszter, 2008). We also note that when using an extension of the model that includes temporal shifts/delays in the temporal modules (as derived in Delis et al, 2014), we did not obtain any task discrimination gain but VAF could be higher. This suggests that the task-related temporal structure of muscle patterns is well explained by the extracted temporal modules and that any minor variations in muscle timings likely require time shifts of those modules. Also it is worth noting that the preprocessing of EMG data tends to filter out measurement noise such that the analyzed signals comprised mostly of physiological within-task and between-task variations whose relative weight was indirectly assessed via the decoding analysis.

In space, the module extraction algorithm grouped muscles into a few (4-7) spatial modules consisting of muscles from different body parts and hemibodies suggesting that groupings did not contain only anatomically coupled or neighboring muscle groups. Also, each spatial module was typically activated to perform many different movements and each movement was performed using the simultaneous activation of many spatial modules (see Figure 6 and Supplementary Material A: modulation of the a coefficients as a function of the initial an final position). This suggests that the spatial modules are not direction-specific but rather functional groups of muscles shared across movements whose weighted recruitment actually codes the task being performed (Tresch et al, 1999; Torres-Oviedo et al, 2006; Delis et al, 2013b, 2014; d’Avella et al, 2006, 2008). The spatial modules were also more variable across participants than the temporal ones. First, this could be due to the fact that subjects had a different optimal number of spatial modules, thus requiring different muscle groupings. Second, such inter-subject differences in muscle synergies have been reported in previous studies (Hug et al, 2010; Guidetti et al, 1996; Frère and Hug, 2012) and are expected to be more pronounced when more muscles are recorded. Indeed, in complex motor tasks (e.g. whole-body-reaching), spatial variability may increase for multiple reasons: different skin conductions or muscle characteristics, different motion kinematics and dynamics, etc. Furthermore, different muscle groupings across individuals may be the outcome of learning or developmental processes, which have recently started to be investigated (Dominici et al, 2011). Therefore, the modular control hypothesis is compatible with the finding that different participants could exhibit different motor modules. However, some muscles such as left tibialis anterior and peroneus appeared to work in synergy for all participants.

### 4.2 VAF and discrimination power compared to other studies

We found that a small set of spatial and temporal modules described muscle activations during performance of a wide range of whole-body pointing movements. This parsimonious representation explained a large part of the between- and within-task variability of the EMG recordings (significantly more than chance), although the actual VAF values we obtained here may be relatively low compared to other studies. One main reason is that the present muscle patterns exhibited a higher level of variability than other studies as a result of a) extracting modules from single-trial data (30 trials for each motion direction), i.e. not resorting to averaging and b) studying a very large set of different tasks (72 distinct movement types, which gives a total of 2160 trials). When extracting modules from trial-averaged data, the average VAF across subjects was 90%, which is comparable to VAF values found in other studies using a smaller number of distinct movements and time-shifts (d’Avella et al, 2006). However, decoding scores could not be evaluated if using averaged data. Note also that our formula for computing the VAF was relatively conservative compared to other formulas that could give arbitrarily larger VAF values, for instance by replacing the mean muscle pattern by zero in the denominator of Eq. 2(Torres-Oviedo et al, 2006). Notwithstanding this, the low VAF and the residual reconstruction error could be taken as evidence that space-by-time modularity can only give a crude description of muscle patterns unless the number of modules is increased. This was already noticed in another study where high VAF levels were needed to accurately reproduce single muscle patterns (Zelik et al, 2014). Here we showed that the low-dimensional EMG description was nevertheless associated to a surprisingly high decoding performance in spite of the fact that task decoding is not an objective of the decomposition algorithm.

Interestingly, the spatial (muscle) dimension of EMG activity appeared to carry more task information than the temporal dimension, which is consistent with previous findings (Delis et al, 2014). The low task information carried by the temporal modules may be partly explained by the fact that muscle signals need to be normalized in time to have equal lengths before being input to matrix factorization algorithms. Time normalization is useful in order to align trials with different durations (and mandatory in current NMF-based methods), however its impact on the task information carried by the resulting signals needs to be investigated further. It is also unlikely that varying the speed instructions (i.e. including different speed conditions in the analysis) would have improved the task discrimination power of the temporal modules as this was not the case for planar arm reaching movements (Delis et al, 2013b). Strikingly, however, our decoding results are comparable and even higher than the ones obtained in the simpler 2-dimensional arm reaching study mentioned above (86% versus 80%) as well as in other studies investigating grasping movements (Weiss and Flanders, 2004; Overduin et al, 2010; Leo et al, 2016). This decoding gain can be explained by two main differences with prior work. First, we used a more flexible model of muscle activation modularity, namely the space-by-time decomposition, which was shown to have higher movement discrimination power compared to alternative models (Delis et al, 2014). Second, we investigated a whole-body reaching task, for which, in contrast to arm reaching and grasping, a) a large number of muscles with complementary functional roles can be recorded using surface EMGs and b) activations of several muscles are expected to be markedly different across tasks. We also note that, in this study, a larger number of temporal and spatial modules was required to achieve these decoding values. This difference indicates that the number of dimensions is dependent on the set of tasks under consideration, hence, to draw more general conclusions about dimensionality, it is important to examine other motor behaviors that are as unconstrained as possible. The task dependency of the modules can be understood if we consider similar reaching movements (e.g. to neighboring targets). In this case, task decoding would be considerably more difficult than discriminating between very different motor behaviors as considered here, where ensemble EMG patterns differ clearly. This point is confirmed via our confusion matrix analyses. Overall, the decoding analysis reveals that recruitment of muscle groupings (spatial modules) is highly dependent on the direction of hand displacement. Their combination with adequate timing signals (temporal modules) via the descending motor commands (activation coefficients) leads to unequivocal characterization of distinct movements.

### 4.3 Space-by-time modularity for motor control

Considering the neural basis of a space-by-time modular scheme, our modeling implicitly assumes that the (invariant) spatial and temporal modules may be stored in certain CNS areas. Then, the (variable) activation coefficients may be triggered by higher-level cortical structures that would generate the descending neural command which would recruit a specific set of modules to coordinate the task-specific recruitment of several muscles and execute the movement at hand. This view is compatible with evidence that, when stimulated, the motor cortex in the primate brain is able to coordinate behaviorally relevant actions, whereby neuronal activity may trigger such goal-directed, multijoint reaching movements (Graziano, 2006). Recently, similar evidence for a high-level encoding of ethological actions has been found in the precentral gyrus of patients undergoing brain surgery (Desmurget et al, 2014). Our findings are not incompatible with such a hierarchical neural implementation of action since the tested modular decomposition was quite effective for task encoding and muscle pattern approximation. However, it is undeniable that the space-by-time decomposition gives only a crude picture of the recorded muscle patterns on single trials. Theoretically, all trial-to-trial EMG variations should be captured by the model (neglecting measurement noise) but a much larger number of modules would be required to achieve VAF above a 90% threshold for instance. This issue was already identified with other existing muscle synergy models for locomotion (Zelik et al, 2014). While we have provided some computational arguments explaining why the VAF is lower than the one observed in prior studies on the topic, we cannot dismiss a potential departure from the proposed feedforward modular control scheme. Indeed, the role of feedback is largely neglected in such models where it might be relevant to better replicate the recorded muscle patterns. Hence we speculate that the high movement discrimination capacity but approximate muscle pattern reconstruction ability may reflect the fact that feedback and/or intermittent control processes occur during motor execution and thus, these processes should be modeled in muscle synergy studies (see below). More generally, the question of falsifying the muscle synergy hypothesis has proved to be difficult although it has been tackled by several neurophysiological and computational works as discussed hereafter.

### 4.4 Critical evaluation of modular motor control

A large number of recent studies attempted to assess modular organizations in terms of their effectiveness in motor control and learning (Kutch and Valero-Cuevas, 2012; Valero-Cuevas et al, 2009; d’Avella and Pai, 2010; Berger et al, 2013; Berger and d’Avella, 2014; Bengoetxea et al, 2014; Inouye and Valero-Cuevas, 2016). In the same vein, other authors have proposed approaches to address the plausibility of modularity in motor control (Giszter, 2015). In particular, monkey electrophysiology (Graziano et al, 2002; Holdefer and Miller, 2002; Overduin et al, 2012, 2014, 2015), human neuroimaging (Asavasopon et al, 2014; Rana et al, 2015) and computational studies (Laine et al, 2015) were employed to investigate the neural origins of motor modularity. Also, modeling studies examined whether optimal motor control can be implemented by modular control schemes (Nori and Frezza, 2005; Chhabra and Jacobs, 2006; Berniker et al, 2009; Neptune et al, 2009; Alessandro et al, 2013a). Finally, studies of human motor behavior investigated the robustness of modules by imposing alterations on muscle coordination of healthy individuals (de Rugy et al, 2012, 2013; Nazarpour et al, 2012a; Steele et al, 2015) and testing muscle activations in clinical populations (Gizzi et al, 2011; Clark et al, 2010; Cheung et al, 2012; Roh et al, 2013).

Here, we tried to approach this issue in agreement with the guidelines developed by Gao and Ganguli (2015) in neuroscience. First, to assess whether modularity may be employed as a strategy for “simplifying” the degrees-of-freedom problem of motor control, modularity should be examined in high-dimensional spaces. In this vein, we propose the design of an experiment that comprises as many movements as possible (with numerous repetitions of the same movement) and involves time-varying EMG recordings of as many muscles as possible (Steele et al, 2013, 2015). Our experiment here comprises more tasks (understand movement types here) than most other studies and also considers a complex daily-life motor behavior of whole-body reaching while standing for which a large interindividual and intra-individual variability may exist (Berret et al, 2009; Hilt et al, 2016). Second, we evaluated the functional role of the model by looking at the extent to which it could represent such a wide variety of muscle patterns with between-task and within-task variabilities. Third, we compared the performance of the modular model with alternative non-modular models in terms of task decoding, data approximation and dimensionality reduction (i.e. the number of parameters to be specified on a single trial). **Here, we could only provide qualitative observations and comparisons with parameters of the non-modular models that were chosen empirically, thus they were not fully optimized. Nevertheless, at equivalent number of parameters, the modular model provided a better description of muscle patterns from both task discrimination and data approximation viewpoints, though more participants and a more thorough search in the parameter space of the EMG signal would be required** to confirm such a tendency. Future work in this direction is needed, in particular to understand if higher VAF could be achieved with more advanced synergy models and if temporal modules that carry more task information could be identified.

### 4.5 Future work

Future research directions should involve investigating alternative formulations of the modular control hypothesis that allow refining motor programs by adapting the modular decompositions for specific task demands, possibly assuming intermittent control (Karniel, 2013). Considering how well the extracted modules allow reconstructing individual muscle activities (Zelik et al, 2014) seems also pertinent to understand if all critical features of muscle activity are considered when generating genuine muscle patterns from a small number of invariant modules. Finally, considering variants of the module extraction algorithm may improve the quality of data approximation and task discrimination. For example, a method that incorporates the task discrimination objective within the module extraction process was shown to identify decompositions with nearly perfect task discrimination power while preserving the same levels of VAF (Delis et al, 2015). Developing methods allowing to avoid time normalization and binning and allowing to consider recruitment of modules via feedback signals would also be very useful especially in order to understand better which neural pathways contribute to the recorded muscle activity and shape the modular organisation we identified here. Of particular interest will be to inform our computational framework with a more detailed account of the underlying physiology of the motor system including the response latencies of different neural pathways as well as the interactions of descending drives and peripheral impulses (Maier et al, 2005; Burke et al, 1992; Darian-Smith et al, 2013; Chakrabarty and Martin, 2011; Galea and Darian-Smith, 1997). In conclusion, further computational and experimental work is required to investigate the motor modularity hypothesis for the neural control of movement.

## Acknowledgements

This research was supported by the « Institut National de la Santé et de la Recherche Médicale » (INSERM), the « Conseil Général de Bourgogne » (France) and the « Fonds européen de développement régional » (FEDER).

